# HIV-Tocky system to visualize proviral expression dynamics

**DOI:** 10.1101/2023.08.10.552733

**Authors:** Omnia Reda, Kazuaki Monde, Kenji Sugata, Akhinur Rahman, Wajihah Sakhor, Samiul Alam Rajib, Sharmin Nahar Sithi, Benjy Jek Yang Tan, Koki Niimura, Chihiro Motozono, Kenji Maeda, Masahiro Ono, Hiroaki Takeuchi, Yorifumi Satou

## Abstract

The stably integrated pool of HIV-1 proviruses in the host genome stands against curative strategies. This reservoir is extremely heterogeneous with respect to host cell type, anatomical location, integration site, and replication fitness. During the initial phase of infection, only a few infected cells can resist host immune clearance or cytopathic effect and establish this resistant pool. The mechanisms underlying HIV latency initiation are not fully resolved yet. In the current study, we propose and validate a new reporter model for monitoring HIV-1 provirus silencing and reactivation using Timer of cell kinetics and activity (Tocky). HIV-Tocky system uses a fluorescent Timer protein whose emission spectrum spontaneously shifts from blue to red to reveal HIV-1 provirus dynamics. We dissected provirus transcriptional phases into early, persistent, recently silenced, and latent. To our knowledge, this is the first report to distinguish two latent subsets: a directly non-expressing population and a recently silenced after brief expression. In-depth integration site analysis suggested that the distribution of proviruses in directly latent cells was similar to that in actively transcribing cell population, whereas recently silenced cells tended to harbor proviruses integrated into heterochromatin. Furthermore, we established a library of various single integration clones at which we utilized to demonstrate the efficiency of the block-and-lock strategy by capturing the fast dynamics of silencing that were overlooked in previous models. In summary, we propose HIV-Tocky system to serve as a time-sensitive model that can capture the dynamics of provirus expression, making it a useful tool for HIV latency research.

**Significance Statement:** Determinants of HIV-1 latency establishment are yet to be elucidated. This reservoir comprises a rare fraction of infected cells that can survive host and virus-mediated killing. *In vitro* reporter models so far offered a feasible means to inspect this population, but with limited capabilities to dissect provirus silencing dynamics. Here, we describe a new HIV reporter model (HIV-Tocky) with dual fluorescence spontaneous shifting to reveal provirus silencing and reactivation dynamics. This unique feature allowed; for the first time, identifying two latent populations: a directly latent, and a recently silenced subset, with the latter having integration features suggestive of stable latency. Our proposed model can help address the heterogeneous nature of HIV reservoirs and offers new possibilities for evaluating eradication strategies.

**Classification:** Biological Sciences, Microbiology.

## Main Text

### Introduction

HIV continues to be an unresolved global public health concern (1) and is one of the three world’s deadliest infections (2). Combined anti-retroviral therapy (cART) showed an incredible effect to turn HIV-1 infection from a life-threatening infectious disease to a chronic viral infection. However, cART is unable to fully eradicate the virus from infected individuals (3) because it is not cytotoxic on virally infected cells and does not block clonal expansion or provirus expression (4).

During the acute phase of HIV-1 infection and prior to cART initiation, most infected cells expressing viral antigens will be eliminated either by host immune clearance or the viral cytopathic effect (CPE) (5). Simultaneously, a reservoir is established through a small pool of stably integrated proviruses in the host genome of infected cells that can escape CPE and immune clearance with the eventual support of host cell survival and clonal expansion (6–8). This reservoir contracts only mildly even after years of antiretroviral therapy (ART) and holds a threatening reactivation capacity once ART is interrupted (3, 9).

HIV-1 reservoir is heterogeneous in terms of reservoir site, provirus intactness, and replication fitness. The reservoir is mainly composed of long-lived resting memory CD4+ T cells, while macrophages, microglia, and dendritic cells can also contribute to specific anatomical sanctuaries (10, 11). Defective proviruses are predominant *in vivo* (12); however, not all intact integrated proviruses are inducible (13). Although this may be partially explained by their stochastic inducibility (12–14), their reactivation dynamics are unclear. Furthermore, integrated proviruses from ART-treated individuals are not uniformly transcriptionally silent, and their individual transcriptional status is governed by interlacing transcriptional, post-transcriptional, and epigenetic mechanisms operating at provirus integration sites (15–18). This convoluted ability of the HIV-1 provirus to hide in latent reservoirs represents the principal obstacle to an HIV-1 cure (3, 19–21).

Interrogation of this integrated reactivatable pool of proviruses has relied on *in vitro* assays for long owing to its rarity. Recent advances in multi-omics analysis have revolutionized the ability to examine this pool in people living with HIV (PLWH) *ex vivo* through which crucial information on the determinants of its maintenance has been obtained. The outstanding key question is how exactly the small pool of reservoir cells is initially selected to survive among these extremely heterogonous infected cells.

Meanwhile, utilizing HIV-1 *in vitro* reporter models has widely aided the advents in eradication strategies. We previously established the widely distributed provirus elimination assay (WIPE assay) as an *in vitro* HIV infection model with hundreds of infected clones that can be maintained for several months (22). This model recapitulated the complexity of infected cells reported for *in vivo* HIV-1 infection, as well as the persistence of defective HIV-1 proviruses.

However, the evaluation of provirus expression using the WIPE assay model requires cell lysis or permeabilization to quantify the viral RNA or protein.

Several generations of dual-fluorescent viruses utilizing green fluorescent protein (GFP) as a reporter for productive infection have been developed. They offered useful information on the early silencing of the HIV-1 provirus shortly after infection (6, 23), the *in vitro* provirus inducibility in response to various latency-reversing agents (LRAs) (24), and addressed the convenience of the shock and kill strategy (25–28). In spite of that, GFP is a stable fluorescent protein with a long half-life (29), which results in a gap between the timing of HIV-1 expression and long-term GFP positivity, which in turn limits the temporal resolution of GFP in capturing provirus dynamics.

To overcome difficulties in analyzing temporal dynamics of transcription *in vivo*, recently a breakthrough was made in immunology by establishing a Timer of cell kinetics and activity (Tocky). This Tocky model uses a fluorescent Timer protein that spontaneously changes its emission spectrum (30, 31). In this study, aiming at the simultaneous visualization of provirus expression dynamics, provirus intactness, and integration landscape, we propose a recombinant HIV construct equipped with this fluorescent Timer protein (HIV-Tocky). We showed that by synchronizing its emission spectrum spontaneous change to provirus expression, Timer fluorescence can punctually trace provirus temporal dynamics from expression to silence. We investigated the integration characteristics of different provirus expression statuses. Moreover, we traced infected cells for clone formation and probed their response to potential latency-promoting agents (LPAs). This *in vitro* model for tracing real-time provirus kinetics provides a paradigm shift in the way latent reservoir composition is described and further assessed for eradication strategies.

## Results

### Establishment of HIV-Tocky system equipped with Fluorescent Timer reporter to capture provirus transcriptional dynamics

Previously, the Ono group developed the Tocky model as a new tool for analyzing transcriptional dynamics using a Fluorescent Timer protein (31), which emission spectrum spontaneously changes from blue to red fluorescence following chromophore maturation (30). As the maturation half-life of the Timer-blue chromophore is 4 h and the mature Timer-red protein is stable with a half-life is 122 h (32), the Tocky system allows the analysis of the dynamics of gene expression and reactivation. Thus, to analyze HIV-1 proviral expression dynamics, we designed a recombinant HIV-1 NL4-3-based molecular clone (33) harboring the coding sequence of Timer protein in the *nef* region, where Timer expression is under the control of the HIV-1 promoter in the 5’ long terminal repeat (LTR); (HIV_Timer_) (Figure 1A; upper panel). The concept of utilizing the temporal shifting feature of Timer fluorescence to visualize HIV-1 proviral expression is best understood by describing the possible fates of the host cell after infection. Infected cells can have one of the following two immediate fates: firstly, an infected cell will actively transcribe mRNAs from the integrated provirus harboring the Fluorescent Timer protein (Timer-FP), rendering it positive for blue fluorescence but still negative for red fluorescence (B+, early). Secondly, infected cells will directly become latent, with no or sub-threshold provirus expression and without any detectable Timer expression (TN). Proviruses within the B+ population can either continue to be transcribed or stop expressing. This creates a mixed population with double positivity for blue and red fluorescence (B+R+, persistent). Here it is considered that most Timer-positive cells will be eliminated by virus-induced CPE and only a small fraction of the B+ or B+R+ cells that have escaped CPE and stopped transcription will eventually lose Timer-Blue fluorescence and acquire the red fluorescence forming the (R+) population. Figure 1B shows a conceptual scheme for the proviral expression dynamics in a single-round infection HIV-Tocky model. To establish a proof-of-concept for the HIV-Tocky system, we infected Jurkat T cells with HIV_Timer_, and analyzed Timer expression dynamics by flow cytometry (Figure 1C). Timer fluorescence was first detectable at 24 h post-infection and evolved in a fan-like movement from B+ to R+, as previously demonstrated by the Nr4a3-Tocky system, at which Timer protein is inducible upon T cell stimulation (30). The R+ population, representing proviruses with recently arrested transcription, started to be evident from 3 days post-infection (dpi) and accumulated gradually over time. Timer-FP positivity peaked at 3 dpi and decreased over time, presumably due to CPE and/or cell cycle arrest induced by viral protein expression in B+ and B+R+ cells (Figure 1E; left panel). Next, in order to identify infected cells in an LTR-independent manner, we modified HIV_Timer_ by inserting the expression cassette of nerve growth factor receptor lacking the intracellular domain (ΔNGFR) in the *nef* region (HIV_TNGFR_, Figure 1A; lower panel). NGFR expression in HIV_TNGFR_ is driven by the constitutively active promoter of *Eukaryotic Translation Elongation Factor 1 Alpha 1(EEF1A1)*(34), and upon provirus integration, NGFR will be expressed on the surface of infected cells enabling the identification of infected cells by flow cytometry irrespective of provirus expression. Infected cells with no proviral expression will be positive only for NGFR but negative for Timer-FP. We infected Jurkat T cells with the pseudotyped single-round HIV_TNGFR_ virus. We analyzed the infectivity and Timer-FP positivity by flow cytometry over time (Figure 1D) and found that the pattern of Timer-FP expression was similar to that of HIV_Timer_ infection and that only a small percentage of infected cells expressed the provirus at all time points, even in the highly proliferating Jurkat T cells (Figure 1E, right panel and Figure 1F), which is consistent with a previous report on primary CD4+T-cells’ infection (25). To evaluate the ability of the HIV-Tocky system to capture provirus dynamics in different cell lineages, we infected THP-1 monocytic cells with the same recombinant viruses. Similar to Jurkat T cells, the Timer-Blue fluorescence appeared 24 h after infection, and R+ cells peaked at 4 dpi (Figure 1G). Timer-FP-expressing cells constituted a small proportion of all infected cells and gradually decreased over time (Figure 1H). To confirm whether Timer expression is correlated with viral gene expression, we sorted each Timer fraction after infecting Jurkat T cells with HIV_TNGFR_ (Figure S1) and quantified viral unspliced and multiply spliced mRNA using reverse transcription polymerase chain reaction (qRT-PCR). The level of early produced and multiply spliced Tat/Rev transcripts was the highest in B+ cells and the lowest in R+ cells. The level of late unspliced Gag transcripts was similar between the B+ and B+R+ populations but low in the R+ population (Figure 1I). These results showed that Timer expression successfully captures viral gene expression dynamics. Furthermore, we analyzed the copy number of cell-associated viral DNA by droplet digital PCR (ddPCR). While there were approximately 1.6 copies of Gag DNA observed per infected cell (Figure S2), 2-LTR DNA, a form of unintegrated viral DNA, was undetectable (data not shown). This suggests that most of the cell-associated viral DNA consisted of integrated proviruses. Collectively, these data indicate the successful establishment of the HIV-Tocky model that can capture the HIV-1 provirus dynamics of reactivation and silencing.

**Fig. 1.**
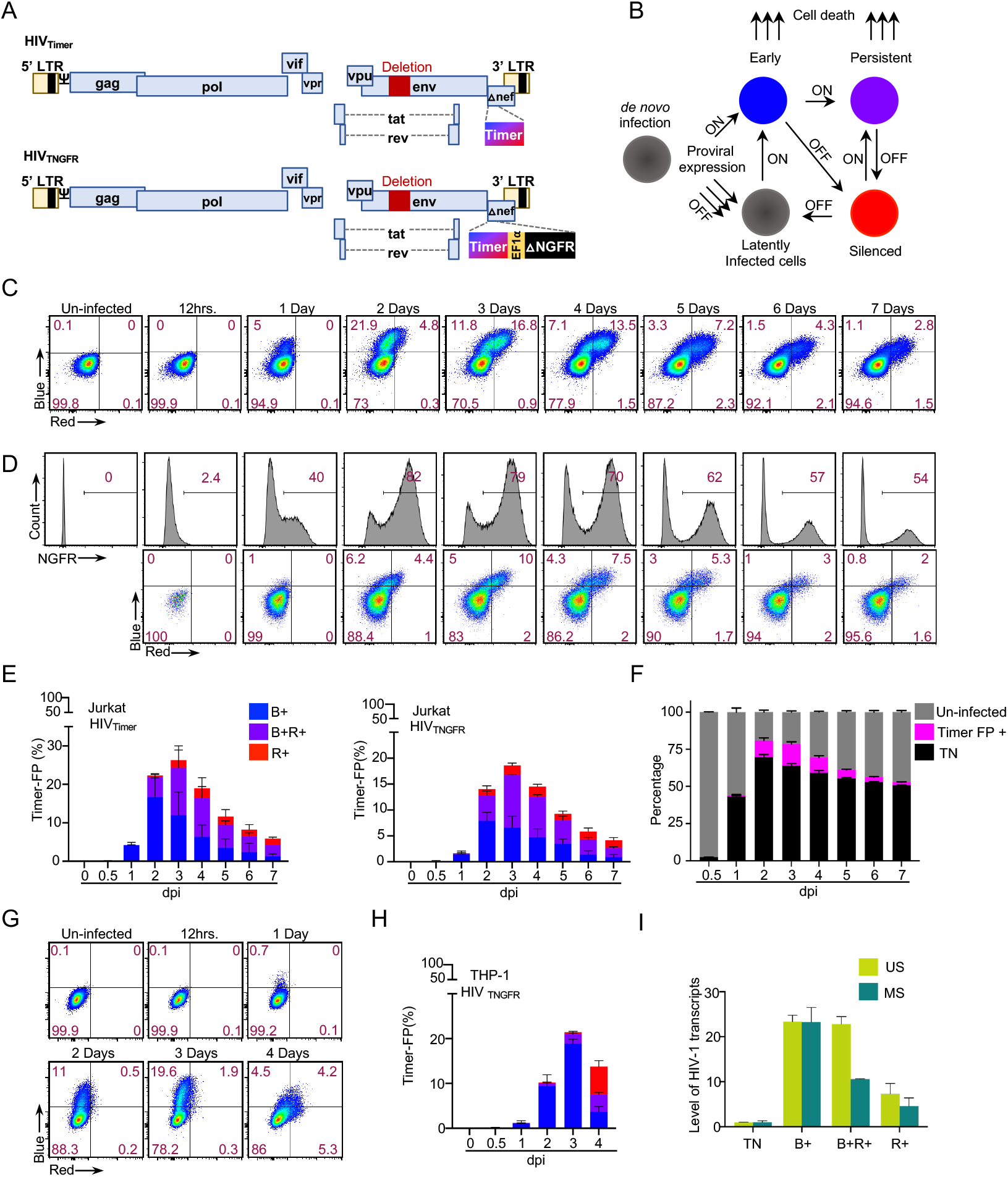
Establishment of *in vitro* HIV-Tocky reporter system to monitor provirus expression dynamics. (A) Schematic representation of HIV-1-Timer constructs (top: HIV_Timer_) and (bottom: HIV_TNGFR_). (B) Schematic for Timer protein expression during different phases of provirus expression or silencing. (C) Representative flow plots from time course infection of HIV_Timer_ in Jurkat T cells. Jurkat cells were infected by HIV_Timer_ virus adding and followed up for Timer expression until day 7 post-infection. The graph is representative of three independent experiments. (D) Jurkat T cells infection by HIV_TNGFR_. Histogram plots representing ▵NGFR percentages until day 7 post-infection (upper panel). Representative Timer-axis shift by flow plots from time course infection (lower panel). The graph is representative of three independent experiments. (E) Representative bar graphs denoting the percentage of each Timer population from infection experiments of Figure 1C in the left panel and of 1D in the right panel. For each graph (n = 3 biologically independent experiments, mean ± SD). (F) Stacked bar graphs representing Timer FP+ fraction (in magenta) in comparison to TN (black) or uninfected cells (grey) across infection time points from 1D. (G) Representative flow plots from time course infection of HIV_TNGFR_ in THP-1 cell line. Cells were infected by HIV_TNGFR_ virus adding and followed up for Timer expression until day 4 post-infection. (H) Representative bar graphs denoting the percentage of each Timer population from THP-1 infection experiment. (n = 3 biologically independent experiments, mean ± SD). (I) Total RNA isolated from each Timer population was subjected to SYBR green RT-qPCR analysis. Unspliced Gag (US) in teal and multiply spliced tat/Rev (MS) in lime green. HIV-1 mRNAs were quantified relative to cellular 18s rRNA. (n = 2 biologically independent experiments, mean ± SD).

### Provirus integration site analysis of Jurkat T cells infected with HIV-Timer virus

The interplay between genetic and epigenetic circumstances at the HIV-1 provirus integration site impacts its fate of either expression or silencing (26, 35–42). Using the HIV-Tocky system, we aimed to analyze the relationship between HIV integration sites (ISs) and categorized provirus transcriptional status by Timer-FP. As described above, the B+ cell population is composed of infected cells with heterogeneous fates, including continuous expression, apoptosis, or “to-be-latent”, whereas the R+ population is homogenously enriched with infected cells harboring the proviruses that were once expressing and later their expression was terminated. Thus, to investigate latency-initiating circumstances, we compared the integration environment between R+ and other Timer populations by sorting TN, B+, B+R+, and R+ populations (Figure S3) and performing IS analysis. HIV ISs were determined by amplifying the junction between the HIV-1 LTR and the flanking host genome using ligation-mediated PCR as described previously (43). We obtained 12,708 unique integration sites, including 4,750, 3,548, 3,919, and 491 ISs in the TN, B+, B+R+, and R+ populations, respectively. Initially, we investigated the distribution of HIV ISs across chromosomes and compared the data from this study with that from previously published reports, including *in vitro* infection (65,924 ISs) and peripheral blood mononuclear cells (PBMCs) isolated from HIV-infected individuals prior to ART initiation (13,142 ISs) (15). First, we observed a similar enrichment of integration in the chromatin of gene-dense chromosomes 17 and 19 in all three datasets (Figure 2A; upper panel). Second, HIV ISs were similarly enriched in the genic region of the host genome in all three datasets (Figure 2B; upper panel). These data indicate that the insertion of the Timer-FP coding sequence in the HIV genome seems to have little impact on the distribution of HIV-ISs compared to previous reports, both *in vitro* and *in vivo*.

**Fig. 2.**
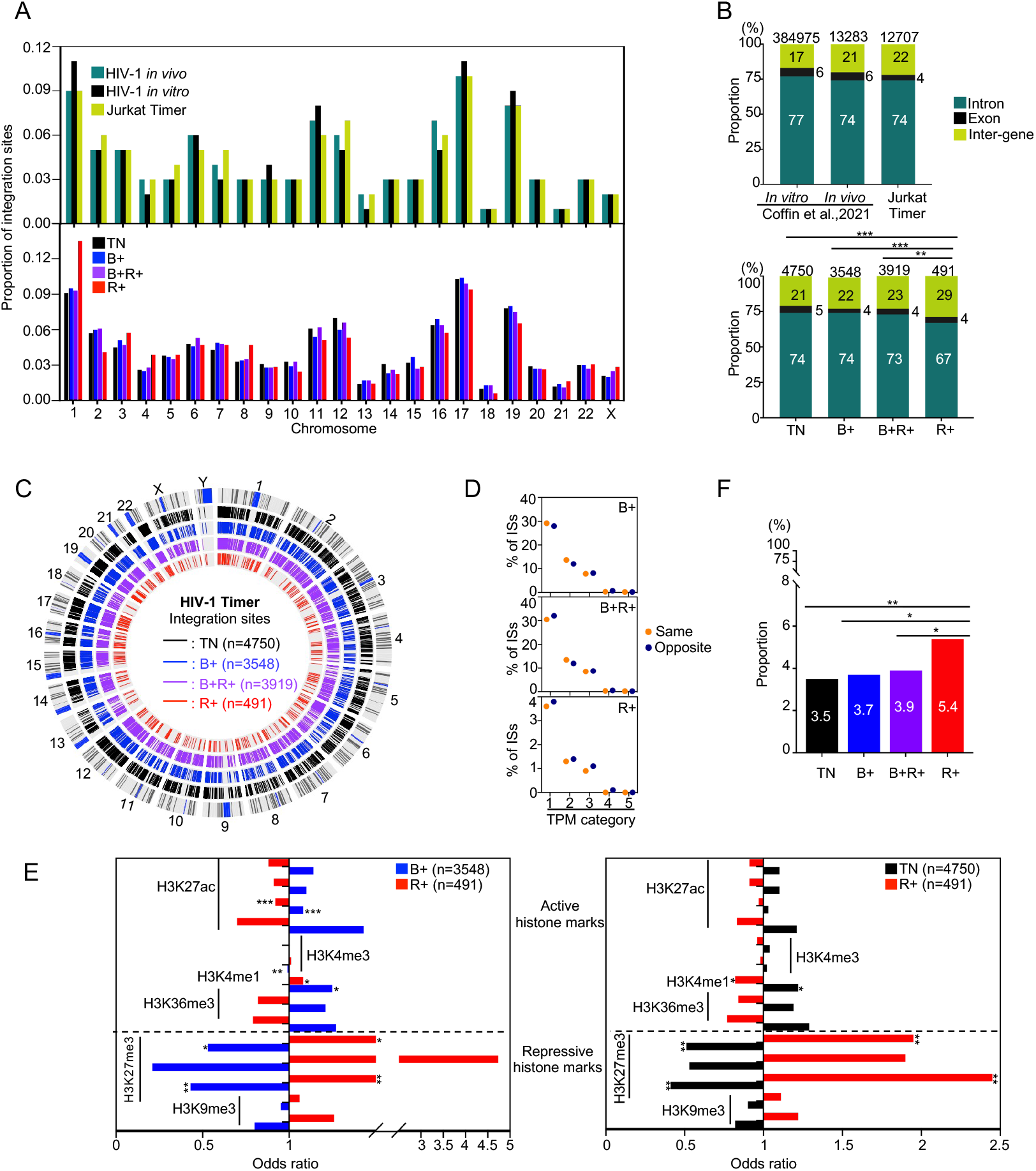
HIV-1 integration landscape in HIV-Tocky *in vitro* system. (A) Relative frequency of integration site distribution in each chromosome. Upper panel: data from Melamed et al., 2022 for HIV-1 *in vivo* and *in vitro* infections are plotted in teal and black respectively in comparison to data from Timer *in vitro* system (in lime-green). Lower panel: Comparing the frequency of integration sites across chromosomes in each sorted Timer population. (B) The frequency of unique ISs within genes or inter-gene distribution in the upper panel: *in vitro* infected PBMCs (2 donors) from Coffin et al., 2021, pre-ART HIV patients from Coffin et al., 2021 in comparison to total detected unique integration sites from Tocky *in vitro* system. Lower panel: The frequency of ISs within genes or inter-genes from different sorted Timer populations. The total number of integration sites analyzed in each fraction is shown at the top of each bar. (C) CIRCOS plot depicting viral ISs across the human genome in Tocky *in vitro* system. Each chromosome is presented on the outer circle and is broken into sequential bins. Blue/grey, black, blue, violet, and red bars indicate G-bands (condensed chromosome region by Giemsa staining for karyotyping), Timer negative, blue, double positive, and red populations, respectively. The number in the parathesis indicates the number of unique ISs detected. (D) Frequency of opposite or same integration across Timer fractions categorized by Transcripts Per Million (TPM); 1:0-50, 2:50-100, 3:100-500, 4:500-1000, and 5:>1000. (E) The odds ratio of viral integration sites within ±2 kb of activating histone marks including H3K27ac, H3K4me3, H3Kme1, and H3K36me3, and the repressive histone marks with H3K27me3 and H3K9me3, comparing B+ and R+ Timer populations (Left panel) and comparing TN and R+ (Right panel). (F) The frequency of unique ISs within ZNF genes was calculated from total genic integrations in each Timer population. For figures (B, E, and F); Statistical significance was assessed by Fischer’s exact test. *<.05, **<.01, ***<.001.

Next, we analyzed the distribution of ISs across Timer-FP populations. Timer-FP ISs were widely distributed across all chromosomes (Figure 2A, lower panel), with no distinct differences across populations. Integrations in the TN, B+, and B+R+ populations favored in gene localization (Figure 2B; lower panel), which agrees to previous reports (7, 44, 45). Notably, the proportion of genic integrations in the R+ population was significantly lower than that in the TN, B+, and B+R+ fractions. Integrations away from the centromere, as per the CIRCOS plot distribution (Figure 2C), were similarly favored by all Timer fractions. HIV-1 integration specifically favors highly expressed gene bodies (46, 47). Therefore, HIV-1 provirus expression is usually studied in the context of the expression level of its host gene, which can have differential yet controversial effects according to the provirus insertion directionality (48, 49). We next obtained public RNA-seq datasets from parent Jurkat T cells to look into the basal expression level of genes with integrated proviruses from our dataset. HIV-1 proviruses in the R+ population tended to integrate into genes with lower basal expression compared to other Timer populations (Figure S4A). Besides, they showed a slight tendency to integrate at longer distances from the transcriptional start site (TSS) (Figure S4B); however, neither of these tendencies was statistically significant. Interestingly, we observed a slightly higher tendency for opposite integration relative to the host gene in the R+ population (Figure S4C) and further traced the tendency of opposite integration across Timer fractions categorized by the basal level of host gene expression. The same direction of integration was preferred in TN and B+ fractions. A slight tendency for opposite integration began to appear in the B+R+ fraction in the lowest expression category and was the dominant tendency in the R+ fraction in all expression categories (Figure 2D).

Next, we investigated whether the integrations in the R+ population were related to distinctive epigenetic features. We obtained published Jurkat uninfected chromatin immunoprecipitation (ChIP) seq datasets (see Materials and Methods for details) and analyzed the frequencies of provirus integration sites within 2 kilobases (kb) of several histone marks by comparing random integrations to B+ and R+ populations (Figure S4D; upper panel). The HIV-1 provirus in both populations favored integration in proximity to active histone marks, but not to repressive ones. This is consistent with previous reports on the integration of HIV-1 provirus (37). Subsequently, we compared the preference for integration near each histone mark between the B+ and R+ populations by calculating the odds ratio. The results showed more frequent integration near the repressive histone marks in the R+ population than in the B+ population (Figure 2E, left panel). We further investigated whether this tendency could be reversed by comparing R+ to the non-expressing subset (TN) and found that TN proviruses showed preferential enrichment close to active histone marks compared to R+ proviruses. (Figure 2E, right panel). HIV-1 integration in genes that encode members of the zinc-finger protein family (ZNF) has been reported to recruit heterochromatin proteins and subsequently support transcriptional repression and long-term persistence (50). Therefore, we sought to identify ZNF integration as an auxiliary epigenetic mechanism controlling latency. We found that approximately 5% of all genic integrations of R+ population occurred in ZNF genes, a significant difference in comparison to any other Timer subset (Figure 2F). Additionally, we observed a similar tendency for ZNF integration between the TN and B+ subsets, and therefore examined their comparative histone enrichment statuses (Figure S4D; lower panel). Enrichment of integrations close to active or repressive histone marks did not show a clear difference between the two populations. These findings indicate unexpected similarities between the TN and B+ subsets, at least at the level of epigenetic control of provirus expression, in contrast to R+ proviruses, which seem to be more strongly influenced by epigenetic control. Collectively, these data suggest the existence of two latent populations, TN+ and R+, with different compositions and/or provirus expression regulatory mechanisms.

### Establishment and integration characterization of HIV-1 Timer clones

Bulk analysis of heterogenous infected cells, each of which has a unique integration environment, provides only the averaged data contributed by all the complex factors governing the establishment and maintenance of latency. To elucidate the clonal heterogeneity of HIV-1 latency, we established HIV-Timer clones, each of which harbored a single copy of the complete provirus. We performed limiting dilution of Jurkat T or THP-1 cells infected with either HIV_Timer_ or HIV_TNGFR_ pseudotyped viruses at 3 dpi (Figure S5). The cloning protocol allows the enrichment of latently infected cells, as infected cells with high viral gene expression are eliminated by CPE. We obtained 17 and 9 Jurkat and THP-1 HIV-Timer clones, respectively. Subsequently, we performed DNA-capture sequencing as previously described (51), which enables comprehensive provirus characterization, including the whole provirus sequence, its respective integration site, and clonal abundance. ISs in established HIV-Timer clones were mostly localized in genic regions (96%) and were widely distributed across human chromosomes (Figure 3A). Host genes carrying the HIV provirus in the HIV-Timer clones showed variable gene expression levels (Figure 3B). Simultaneously, we analyzed the proviral expression status of each clone using Timer-FP. HIV-Timer clones showed wide intra- and inter-clonal heterogeneity in terms of provirus expression, as shown in some representative results (Figure 3C; upper panel). Notably, we did not observe a significant correlation between the basal host gene and proviral expression levels (Figure 3D). We further examined the induction of provirus expression by T cell stimulants or TNF-α (Figure 3C; lower panel and Table 1) and obtained available RNA-seq and ChIP-seq data for activating and repressing histone marks of parent Jurkat T cells to obtain a close-in view of the pre-integration environment in two representative examples of HIV-Timer clones: JO#10 and JO#19. Clone JO#10 had a provirus integrated in an active transcribing gene (*HNRNPM*) (Figure S6A; left panel) while showing a relatively low basal Timer expression (Figure 3C). In contrast, in clone JO#19, we observed a high basal expression for a provirus integrated into a silenced gene (*TSBP1*) (Figure S6A; right panel). Pre-integration histone mark enrichment and HI-C compartment information (52) confirmed the basal transcriptional activity of the host gene in each clone (Figure S6A, S6B, and Table 1). We further investigated whether provirus integration altered the transcriptional activity of the host genes by performing RNA-seq analysis of these two HIV-Timer clones. We did not observe any significant change in the local transcriptome of the host gene after HIV integration in JO#10 (Figure 3E, left panel) or JO#19 (Figure 3E, right panel). Lastly, we quantified the transcription level using the transcripts per million (TPM) value and found a modest increase in the transcriptional activity after provirus integration (Figure 3F and 3G). Notably, the proviruses were integrated in the opposite direction in JO#10 and in the same direction as the host gene in JO#19. Contradicting provirus expression levels can be partially explained by the transcriptional interference phenomenon (48); however, further study of the integration environment of each Timer fraction within a clone can provide valuable information on latency maintenance determinants. Collectively, these data demonstrated the successful establishment of a series of HIV-1 latent clones with heterogeneous provirus expression and variable integration into host genomic regions. Moreover, we were able to concomitantly characterize the whole proviral sequence and proviral expression at single-cell resolution, thereby offering a valuable resource for further investigation of HIV-1 latency mechanisms.

**Fig. 3.**
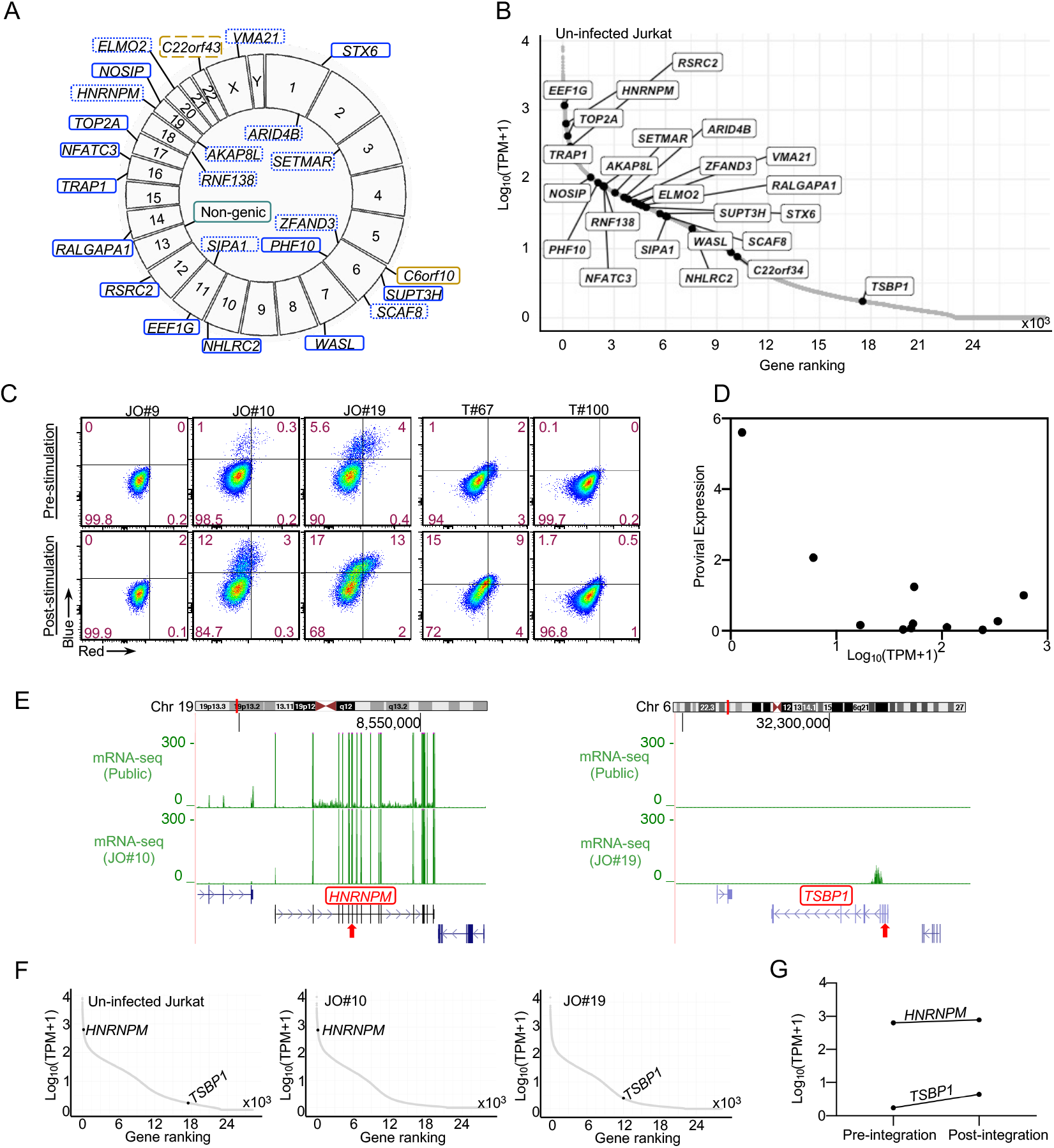
Establishment and characterization of HIV-1 Timer clones. (A) CIRCOS plot demonstrating chromosomal integration site positioning of Jurkat Timer clones for retrieved near full-length proviruses from a total of 17 clones. Color and line coding indicate genic/non-genic positioning, the orientation of integrated proviruses relative to host gene, and gene transcription status in Jurkat cells. Active gene is squared in blue, silenced gene in gold. Solid line indicates the same orientation, and the dotted line indicates the opposite orientation to the host gene. (B) Ranking plot showing TPM values and ranking of each gene of integration of Timer clones are shown from RNA-seq data of uninfected Jurkat T-cell line (pre-integration data). (C) Flow cytometry plots showing Timer FP expression denoting provirus expression in selected single integration Timer clones. Jurkat Timer clones’ basal expression (upper left panel); provirus expression after T-cell stimulation for 24 h (lower left panel); THP-1 clones’ basal expression (upper right panel) and THP-1 provirus expression after TNF-⍺ stimulation for 24 h (lower right panel). (D) Correlation scatter plot of proviral expression percentages from single integration Jurkat Timer clones and level of expression of the corresponding gene of integration. (E) RNA-seq data of Jurkat uninfected T-cell line and RNA-seq data obtained from infected clones JO#10 (right panel) and JO#19 (left panel) are plotted. The position of the integrated provirus in each clone is indicated by a red arrow. (F) TPM values and ranking of each gene of integration of clone JO#10 and JO#19 are shown from RNA-seq data of uninfected Jurkat T-cell line (pre-integration, left panel) and after integration (middle panel; JO#10) and (right panel; JO#19). The observed change in TPM value pre- and post-integration within each clone is demonstrated in (G).

**Table 1.**
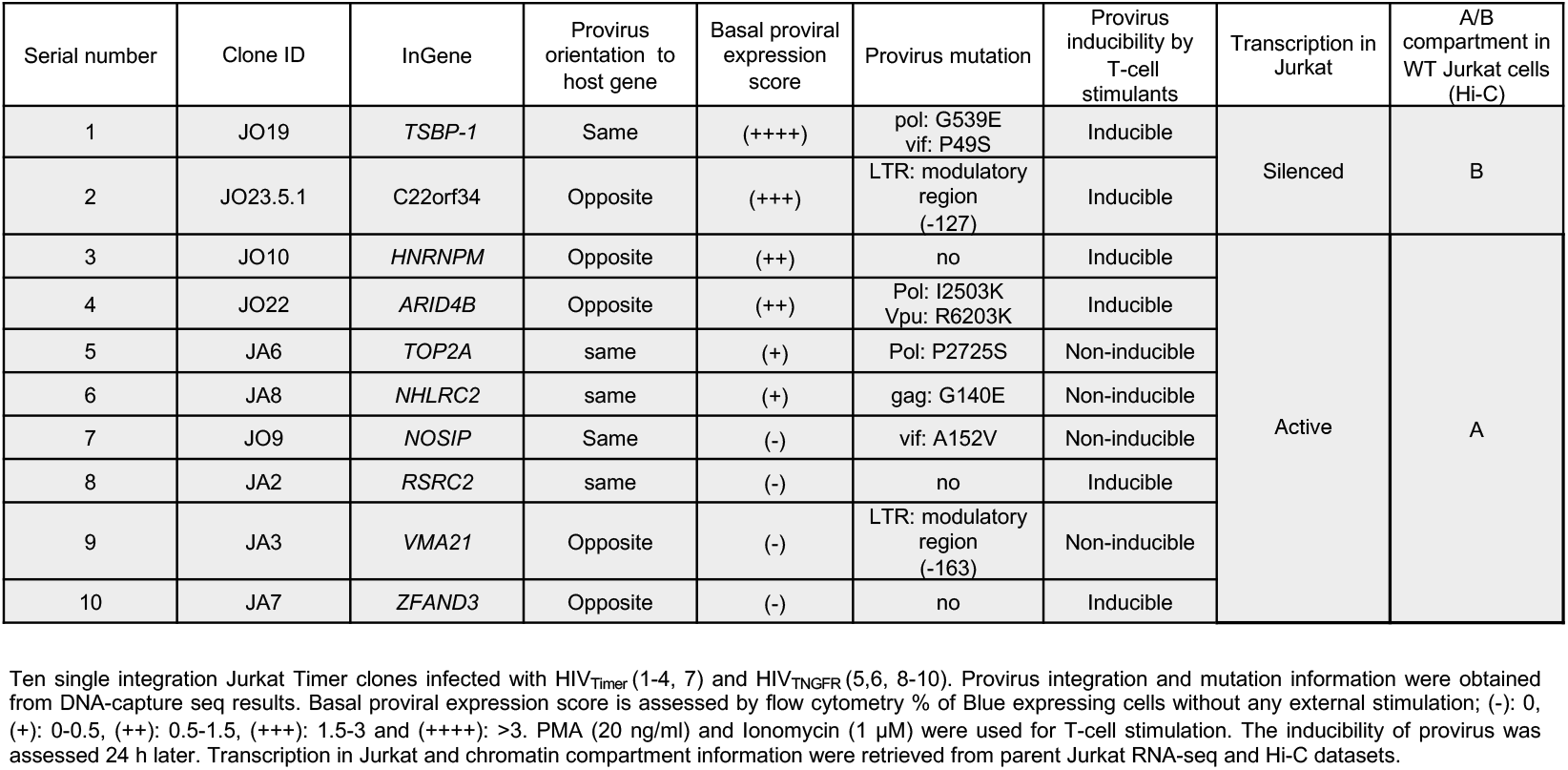
Characterization for established HIV-1-Timer Jurkat clones.

### Application of HIV-Timer clones for the evaluation of Latency promoting agents

One possible application of HIV-Timer clones is in drug screening for latency-reversing or- promoting compounds. Although several latent cell models harboring fluorescent proteins currently exist for monitoring proviral expression, HIV-Timer clones may have some advantages because of their unique characteristic of spontaneous transition from blue to red fluorescence upon provirus transcriptional arrest. Since the maturation half-life of Timer Blue transition to red fluorescence is 4 h, we can detect reactivated or recently silenced proviruses by Timer fluorescence in a highly sensitive manner. Thus, we aimed to investigate the potential of the HIV-Timer system to capture changes in the provirus transcriptional status when challenged with LPAs. First, we used J-Lat 10.6 cells to evaluate LPA activity as a representative of the conventional GFP model. In J-Lat 10.6 cells, a GFP coding sequence is inserted in the *nef* region to monitor proviral expression. Subsequently, we treated the cells with TNF-α to reactivate proviral transcription before adding the LPA reagent. Forty-eight hours after TNF-α stimulation, we treated cells with LPA or dimethyl sulfoxide (DMSO) for an additional 12 h (Figure 4A). We used 2 classes of LPAs: Levosimendan, which promotes Tat degradation, and Triptolide, which inhibits NF-κB activity. We used three drug concentrations according to previous reports (53, 54), and performed a cell viability assay to exclude any non-specific cytotoxic effects on the result. No obvious toxicity was observed at all tested concentrations (Figure S7). We did not observe any change in GFP positivity during the initial 12 h after drug treatment (Figure 4B,4C, and S8). Next, we treated the HIV-Timer clone JO#19 with the same LPAs or DMSO using the same conditions as those for J-Lat 10.6 cells (Figure 4A) and found a significant decrease of B+ cells and a significant increase of R+ cells (Figure 4D) with both LPAs used in comparison to DMSO treatment. Timer transitions in response to Levosimendan or Triptolide are summarized in Figure 4E. Taken together, we demonstrated that the HIV-Tocky system can capture proviral transcription dynamics under latency-promoting effect in a highly sensitive manner compared with a conventional GFP latent infection model.

**Fig. 4.**
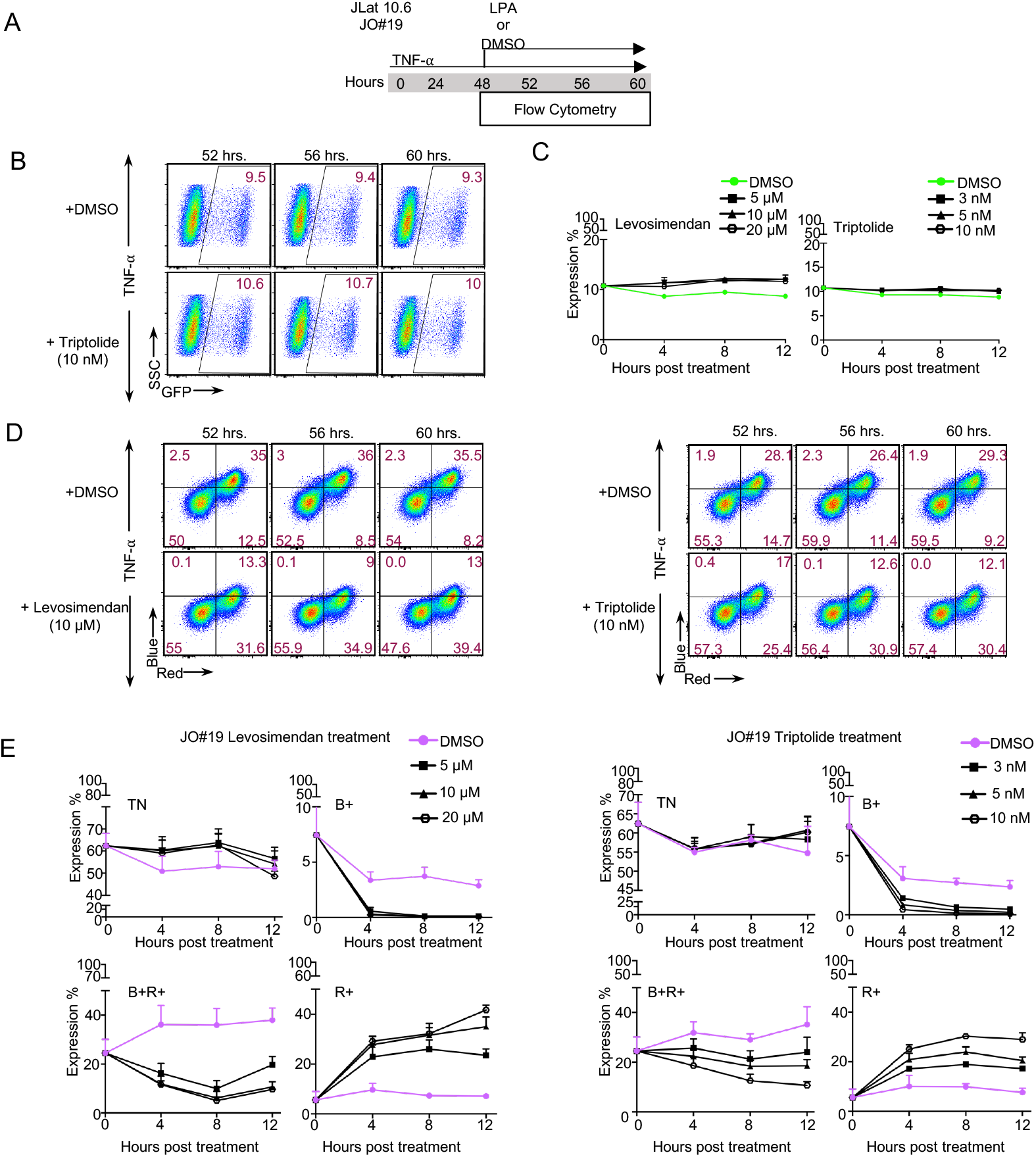
HIV-1 Timer clones as a model for estimating LPAs efficacy. LPAs effect on provirus silencing monitored by Timer transitions or GFP. (A) Assay overview. Schematic of experimental procedure involving drug treatment of Jurkat Timer clone JO#19 or JLat 10.6. Briefly, Jurkat Clone JO#19 or JLat 10.6 cells were stimulated with TNF-⍺ (10 ng/µL) for 48 h, an LPA drug or DMSO was further added for 12 h, and the changes in Timer expression or GFP were monitored, respectively. (B) JLat 10.6 cells were stimulated with TNF-⍺ (10 ng/µL) for 48 h, an LPA drug or DMSO was further added for 12 h, and the changes in GFP expression were monitored. Flow plots demonstrating GFP transitions under Triptolide (10 nM) treatment are shown. Flow plots representing Levosimendan (10 µM) treatment are shown in supplementary figure 8. (C) The expression percentage of GFP is plotted against time under Levosimendan treatment (left panel) or Triptolide (right panel) with 3 stated different concentrations in comparison to DMSO treatment. (n = 3 biologically independent experiments, mean ± SD). (D) Flow plots demonstrating Timer transitions over 12 h treatment with DMSO (upper left panel), Levosimendan (10 µM) (lower left), or DMSO (upper right), Triptolide (10 nM) (lower right panel). (E) Transitions in each Timer population from zero point until 12 h post-treatment are graphed by expression percentage obtained from flow cytometry analysis of each Timer population against time under Levosimendan treatment (left panel) or Triptolide (right panel) with 3 stated different concentrations in comparison to DMSO treatment. (n = 3 biologically independent experiments, mean ± SD).

## Discussion

*In vitro* HIV-1 reporter systems have advanced HIV-1 latency research in several respects. HIV-1 establishes its reservoir at an early stage of infection *in vivo* (21, 55–58). However, elucidating the mechanisms of latency establishment necessitates marking very rare latent infected cells (59) which in turn requires the use of several *in vitro* models that could recapitulate the situation *in vivo* (22, 26, 37, 60). Despite recent revolutionary advances in scrutinizing latent reservoirs *ex vivo* at the single-cell level (16, 61–63), the most recent report by Sun et al. concluded the inability to identify a single universal marker for latently infected cells (64). This signifies the need to improve HIV-1 reporter models to obtain guiding information regarding the obscure latent behavior of HIV-1. The development of the HIV-1-GFP reporter (65) and its use for cell line infection (26, 66) has paved the way for later marking and separation of infected cells from purely latent cells in dual-reporter systems (23, 24). A GFP-modified reporter (HIV_GKO_) facilitated better separation of latently infected cells from infected primary CD4^+^ T cells *in vitro* and provided additional evidence on the reactivation heterogeneity of the latent population (25) reported previously (13, 38). Notably, all the aforementioned models used stable GFP—which has a long half-life (t ½ = ∼ 56 h)—as a reporter (29). This means that some expressing proviruses might lose their expression capacity at a certain point; however, this will not be reflected on the status of GFP positivity because its decay time has not yet been reached. Pearson et al. (67) used a modified short half-life d2GFP (t ½ = ∼ 3.6 h) to provide a more effective measurement of provirus transcription rates. Additionally, Venus fluorescence under LTR control (68) has been used to analyze the dynamic regulation of LTR activity. Both systems demonstrated provirus reactivation and silencing kinetics in a time-discernible manner; however, any temporal heterogeneity of silencing or stochastic re-expression can’t be further tackled. Here, we propose and validate a new HIV-1 reporter system for monitoring proviral expression dynamics with higher temporal resolution than conventional GFP systems by introducing a Timer-FP into a recombinant HIV-1-derived vector (HIV-Tocky) (Figure 1A, B). *In vivo*, the risk of viral rebound is maintained by a rare minority of cells harboring intact latent proviruses (12) (13). In this study, we used env-defective single-infection-cycle HIV molecular constructs to limit the appearance of defective proviruses. The HIV-Tocky system captured the dynamics of the very early phase of HIV-1 infection, including the simultaneous productive infection and latent reservoir establishment as reported *in vivo* (59), in a time-sensible manner in both Jurkat T and THP-1 myeloid cell lines (Figure 1 C, G).

In the HIV-Tocky system, the rare surviving cells that are expected to contribute to early reservoir seeding are represented by the scarce R+ fraction that survived and lost expression before joining the initially silent fraction (TN) (19, 21). To rule out the possibility that the monitored kinetics were affected by the selective growth advantage of the uninfected cells, we additionally constructed HIV_TNGFR_ with an expression cassette for NGFR, which enabled us to distinguish purely latent from uninfected cells. After the selection of infected cells, the same provirus dynamics as in the HIV_Timer_ were replicated (Figure 1D). To the best of our knowledge, the current study is the first to successfully identify a latent cell subset with integrated proviruses that showed transient expression, which was subsequently terminated (R+). Using bulk integration site analysis, we first verified the generation of heterogeneous clones in our HIV-Tocky *in vitro* model as previously reported in our WIPE assay model (22). R+ proviruses demonstrated distinct integration features compared with other Timer populations. They showed a tendency to integrate more in non-genic regions and with orientations opposite to those of the host genes. Genic integration in ZNF-encoding genes was significantly higher in the R+ population. R+ proviruses were more likely integrated in close proximity to transcriptionally repressive chromatin regions. These findings are consistent with some reported attributes suggestive of “deep latency” in intact proviruses retrieved from elite controllers (69) or individuals with HIV under prolonged ART (70), which are shielded from host immune selection. However, we observed such features in a population of proviruses that lost their expression just a few hours after infection, and apart from long-term immune selection forces. HIV-1 latency at the transcriptional level is governed by proviral intactness, host cell factors, transcriptional interference, and epigenetic modifications. In contrast, stochastic provirus expression as mediated by the Tat-feedback loop was reported to control infected cell fate regardless of host cell status or chromatin organization (71, 72). In our dataset, as explained above, the criteria suggestive of “stable” latency were more significant in the R+ proviruses than in TN proviruses, which are proviruses with a decided lack of expression since the onset of infection. A striking finding was the genetic and epigenetic similarities existing between TN and B+ integrations. Given these observations, we can assume that TN population has a heterogeneous composition of proviruses that are more readily able to stochastically turn on expression when aided by the Tat feedback loop or host cell factors. Conversely, R+ is a more controlled population with determined integration features into a quiescent destiny, guided by heterochromatin insertion or opposite directionality. Taken together, our data describe two different layers of latency with differential regulatory determinants at which HIV-1 provirus reactivation and silencing appear to be asymmetric phenomena, with the former being more stochastic and the latter being more controlled.

Next, we generated a library of Timer single-integration clones that could help in future investigations of latency determinants. Timer clones showed a wide variety of integration localities and basal target gene expression. In our model, in which Tat expression and basal LTR promoter activity were intimately coupled, the intra- and inter-clonal heterogeneity of provirus expression agreed with what has been previously monitored (40). After ruling out cell environment differences in a uniform cell line, as well as the role of provirus mutations by confirming Tat and TAR intactness in our clones, we examined the pre-integration environment as a possible determinant of clonal heterogeneity. Our clones showed a preference to integrate in actively transcribed genes (eight out of ten) following HIV-1 preferred integration pattern but in contrast to (26); where they have enriched for latently infected cells prior to clone generation. A detailed mechanistic analysis of the expression pattern of each clone was not performed in the present study; however, the behavior of the Timer clones, despite their limited number, shares some features with previous *in vivo* or *ex vivo* reports. For instance, intact non-inducible proviruses (13) and active expression of persistent proviruses in ART-treated PLWH (73). One notable finding was the high expression of the provirus in two out of ten clones (Table 1, Figure S6A; right panel), despite integration in the heterochromatin region. The factors governing latency reversal are complex. The dynamics of provirus transactivation by Tat have been thoroughly studied and proven to be stochastic (74). The Tat-positive feedback loop shows a robust effect that could determine the fate of provirus expression regardless of cell-state changes (72, 75). Moreover, the Tat feedback loop allowed the induction of expression even in proviruses integrated near human endogenous retroviruses (HERV) LTRs (71), which are known to localize in heterochromatin regions (76). Although this may partially explain the phenomenon, extended experiments by sorting and exploring the integration environment of each Timer population within each clone can provide more clues regarding the molecular mechanisms of provirus silencing and expression. Overall, the available set of the Timer clones demonstrated a reasonable variety of integration and provirus expression and can be used as a novel tool for investigating the mechanisms of latency establishment and reversal aided by the high-resolution tracing of provirus kinetics by Timer fluorescence dynamics.

The ideal HIV treatment should result in remission and eradication. Assuming the uniformity of HIV-1 latent reservoir, the “shock and kill” cure strategy aims to induce provirus expression to facilitate their immune clearance or virus cytopathy. Despite the widespread advocacy of this strategy and hundreds of LRAs being screened (77), none have been reported to achieve the desired reductions (78) in latent reservoirs to secure a non-rebound cure (25). This inefficiency was later attributed to the heterogeneity of the latent reservoir (79), with less than 1% of the latent reservoir being reported to fuel the viral rebound (78). Moreover, owing to the stochastic fluctuations in Tat expression, the inducibility rates and efficiency of LRAs *in vivo* are highly variable (13, 71). Additionally, the ability of some transcriptionally active proviruses to survive *in vivo* and resist immune clearance adds to the challenges facing this strategy. “Block and lock” is an alternative strategy for achieving HIV-1 functional cure that entails enhancing the provirus latent state and getting the provirus promotor into deep and irreversible latency via epigenetic modifications, resulting in a drug-free non-remission status. The concept of provirus “deep latency” has been described to be naturally occurring in 2 rare types of PLWH: elite and post-treatment controllers. This proposes the block-and-lock strategy as a more promising *in vivo* eradication approach. Of note, the fluctuating nature of the Tat positive feedback loop is a fundamental “to-control” target to ensure the full success of the block and lock strategy. In the current study, we demonstrated the feasibility of Timer fluorescence transitions to reliably capture the provirus silencing kinetics which was not possible by GFP (Figure 4) using two LPAs with different modes of action on Tat transactivation. In either case, GFP dynamics remained stable, while the shut-off of provirus expression was clearly represented by the increasing percentage of the R+ population over the same drug exposure time in the Tocky model. Therefore, we demonstrated the ability of the Timer-FP to capture the rapid dynamics of silencing by targeting the Tat-LTR axis. Further experiments on epigenetic changes at the IS, especially in the R+ fraction, can elucidate more on the mechanisms of irreversible provirus silencing.

In conclusion, we propose a novel reporter system for HIV-1 provirus expression dynamics that enables the analysis of the multilayered nature of HIV-1 latency by revealing real-time dynamics of provirus expression. Future studies with more detailed downstream analyses of these dynamic fractions can provide insights into the nature and determinants of their behavior. We propose the HIV-Tocky model as a useful tool for drug screening aided by high-throughput flow cytometry, particularly for modulators of HIV-1 expression fluctuations, which can enhance irreversible latency. Additionally, we showed that the HIV-Tocky system recapitulated the expected dynamics in cell lines of different lineages. Different cellular reservoirs deal differently with HIV-1 integration and latency For example, in monocyte-derived macrophages, proviruses have less preference for integration in actively transcribing genes (80). Future applications in primary myeloid and lymphoid cells may provide more information on latency dynamics and determinants in different tissues and anatomical reservoirs.

## Limitations of the study

This study has some limitations. First, we utilized an *in vitro* model of provirus kinetics that lacked the immune selection effect and relied solely on the virus CPE for clearance. Furthermore, we used cell lines with known high proliferative capacity, which can mask some aspects of provirus dynamics. Thus, further analysis using primary CD4+ T cells or *in vivo* models is required for a more accurate visualization and assessment of HIV-1 provirus kinetics.

## Materials and Methods

### Cell culture

HEK293T cells (human embryonic kidney cell line; American Type Culture Collection (ATCC)) were used for virus production. Jurkat T cells (E6.1 from ATCC) and THP-1 cells (a human monocytic cell line; kindly provided by Dr. Hiroaki Takeuchi [Tokyo Medical and Dental University] were used for infection. JLat10.6 cells were obtained from the National Institutes of Health AIDs reagent Program. HEK293T cells were maintained in Gibco Dulbecco’s Minimal Essential Medium (Gibco, Thermo Fisher Scientific), and suspension cell lines were cultured in Roswell Park Memorial Institute Medium (RPMI 1640; Gibco, Thermo Fischer Scientific). All culture media were supplemented with 10% heat-inactivated fetal bovine serum, penicillin (100 U/ml), and streptomycin (100 μg/ml). All cell lines were maintained at 37°C, 5% CO_2_. Suspension cells were kept in T25 flasks and passaged twice a week, with a cell to fresh medium dilution ratio of 1:3.

### Plasmid construction

We used two recombinant viruses, pNL4-3/ΔBglII/Timer (HIV_Timer_) and pNL4-3/ΔBglII/Timer-EF1α-1′NGFR (HIV_TNGFR_). Both plasmids were derived from pNL4-3/ΔBglII (81). The *env* gene was removed by BglII (82). The *nef* gene between *env* gene and XhoI restriction site was replaced with Timer or Timer-EF1χξ-1′NGFR gene.

### Virus production and infection

Pseudotyped HIV_Timer_ and HIV_TNGFR_ virus stocks were generated by transient co-transfection of HEK293T cells with a plasmid encoding either HIV_Timer_ or HIV_TNGFR_ and VSV-G using Lipofectamine 3000 transfection reagent (Invitrogen, Cat# L3000-015) according to the manufacturer’s protocol. At 28–40 h post-transfection, supernatants were harvested and filtered through a 0.45-μm-pore-size filter to clear cell debris. The virus was precipitated by centrifugation at 13200 ×*g* for 60 minutes at 4°C. Concentrated virions were resuspended in complete RPMI 1640 medium, aliquoted, and stored at -80°C. The amount of p24 Gag in the concentrated virus was quantified using an HIV-1 p24 antigen enzyme-linked immunosorbent assay (ELISA) kit (ZeptoMetrix, Cat# 0801002) according to the manufacturer’s instructions. Jurkat T and THP-1 cell lines’ infection with HIV_Timer_ or HIV_TNGFR_ molecular constructs was performed by cells’ incubation with the viral supernatant diluted in complete RPMI 1640 medium at 10 ng of p24 per 5×10^5^ cells. On assigned sampling time points (12 h until day 7), one third medium change was performed. Unless otherwise specified, the final virus concentration used for infection was equivalent to 10 ng, as titrated by p24 ELISA. All HIV-1 infection and infected cell lines were handled in bio-containment level 3 rooms at Kumamoto University.

### Establishment of Timer clones

For Jurkat Timer clones, cells infected with HIV_Timer_ at 3 dpi were cloned by limiting dilution by seeding 0.3 cells per well in 96 cell culture plates. After approximately 3 weeks, 24 clones were obtained. We confirmed 11 clones for infection by qPCR for *Gag* region (data not shown). THP-1 clones were infected as described above in the virus production and infection subsection. To select the infected cells, NGFR MACS separation was performed according to the manufacturer’s instructions. First, the infected cells were stained with a biotinylated anti-human CD271 (NGFR) antibody (BioLegend, Cat# 345122). Subsequently, the cells were magnetically labelled with Streptavidin MicroBeads (Miltenyi Biotec, Cat# 130-048-102). The cell suspension was then loaded onto a MACS column, which was placed in the magnetic field of a MACS separator. Magnetically labelled NGFR+ cells were retained in the column and further eluted as the positively selected cell fraction after magnetic removal. NGFR+ cells were subjected to limiting dilution, as described for Jurkat clones. Twelve THP-1 clones were obtained. Additional 6 Jurkat Timer clones were generated using the NGFR+ selection method after infection.

### Timer and JLat clones’ drug treatment

To check provirus inducibility in Timer clones, Jurkat Timer clones were treated for 24 h with a combination of T-cell stimulants: phorbol 12-myristate 13-acetate (SIGMA, Cat# P1585) at a final concentration of 20 ng/ml and ionomycin (Nacalai Tesque, Cat# 19444-91) at 1 µM final concentration. For THP-1 Timer clones, recombinant human TNF-alpha (hTNF-⍺, PEPROTECH, Cat# 300-01A) at 10 ng/ml was used for 24 h. In latency-promoting experiments (Figure 4), Timer or JLat 10.6 clones were seeded at 5×10^4^ cells per 12 well plate and treated with hTNF-⍺ at 10 ng/ml concentration. At 48 h, LPA or DMSO was added, and the cells were aliquoted at 4, 8, or 12 h for flow cytometric analysis. The following LPAs were used: Levosimendan (TGI, Cat# L0320) at 5, 10, or 20 µM final concentration and Triptolide (Cayman, Cat# 11973-1MG) at 3, 5, or 10 nM final concentration. Cell viability was assessed using cell counting kit-8 assay (Cat# GK10001). The JO#19 cells were serially diluted and seeded according to the manufacturer’s instructions. Treatment with LPA or DMSO was continued for 8 or 24 h prior to the addition of cell-counting kit-8 reagent. After 4 h of incubation, color absorbance from each well was measured using SPECTRAmax 190 microplate spectrophotometer (Molecular Devices, USA) using a 405 nm filter. Obtained absorbance was further analyzed using SoftMAx Pro Microplate analysis software (version 5, Molecular devices, California, USA). Absorbance was normalized to the no-cell control, and the final cell survival percentage was divided by the DMSO control survival percentage and plotted (Figure S7).

### Timer flow cytometric analysis and cell sorting

For integration site analysis, Jurkat T cells were infected with HIV_Timer_ virus construct as described above. Infected cells (5×10^5^) were aliquoted at 12 h, and 1–7 dpi and cryopreserved. Time-course flow cytometric analysis was performed by thawing and analyzing all time points in the same setting. For Timer-only fluorescence detection, cells were revived in pre-warmed complete RPMI 1640 medium, washed once with PBS, and incubated in near-IR fluorescent live/dead fixable dye (Thermo Fischer, Cat# L34972) to stain dead cells at a 1:1000 dilution in PBS for 30 min at 4 °C. Cells were then washed once and then fixed with 1% PFA in PBS for 15 min at 4 °C in the dark prior to data acquisition. To determine the surface NGFR expression along with Timer, revived cells were first stained for dead cells as described above, and after a single wash in PBS, they were stained with APC-conjugated anti-human CD271 (NGFR) antibody (BioLegend, Cat# 345108) at 1:30 final concentration for 30 min at 4 °C. Cells were washed twice with PBS and fixed in 1% PFA in PBS for 15 min in dark at 4 °C prior to data acquisition. Timer and ΔNGFR fluorescence were measured and/or sorted using SH800 sorter (Sony). Timer Blue fluorescence was detected in the FL1 channel (450/50 nm) excited by a 405-nm laser, while the red fluorescence was detected in the FL3 channel (617/30) excited by a 561-nm laser. Data were analyzed using FlowJo 10.7.1 software (Tree Star, Inc.). For viral mRNA quantification, Jurkat T cells were infected with HIV_TNGFR_ virus construct as described above. The infected cells were collected from day one to 7 post-infection and cryopreserved. Aliquots were then resuspended in warm complete RPMI 1640 medium, washed once with PBS, and stained with near-IR fluorescent live/dead fixable dye (Thermo Fischer, Cat# L34972), followed by staining with APC-conjugated anti-human CD271 (NGFR) antibody (BioLegend, Cat # 345108) at 1:30 final concentration as described above. Cells were finally washed twice with PBS and resuspended in 1% FCS in PBS buffer. The Timer populations were sorted using BD FACSAria III machine at the biocontainment level 3 facility at Kumamoto University. To obtain enough cells for downstream analysis, TN and R+ cells were sorted from day one and day four post-infection, respectively. B+ and B+R+ cells were sorted on days 2 and day 3 post-infection samples.

### qRT PCR for HIV transcripts quantification

RNA (Figure 1I) was extracted RNeasy Mini Kit (Qiagen, Valencia, CA, USA) according to the manufacturer’s instructions with DNaseI treatment. cDNA was synthesized using ReverTra Ace® qPCR RT Master Mix (Toyobo, Osaka, Japan, Cat#FSQ-201) according to the manufacturer’s instructions. Relative cellular HIV mRNA levels were quantified using SYBR qPCR assay using primers to detect unspliced *Gag* mRNA as described in (Douek et al, Nature, 2002) and multiple spliced *tat/rev* transcripts; 5’ for-TCA GAC TCA TCA AGC TTC TCT ATC AAA GC-3’ and 5’ rev-GAT CTG TCT CTG TCT CTC TCT CCA CC-3.’ Viral transcripts were normalized to 18S rRNA. The assay was performed using an Applied Biosystems StepOnePlus Real-Time PCR System (Thermo Fisher Scientific). Relative cell-associated HIV mRNA copy numbers were determined in a reaction volume of 20 µL with 10 µL of Thunderbird SYBR qPCR mix (Toyobo, Osaka, Japan Cat#QPX-201T), 0.3 µM of each primer and 2 µL of cDNA. Cycling conditions were 95 °C for 1 min, then 40 cycles of 95 °C for 15 s and 60 °C for 1 min. Real-time PCR was performed in triplicate and relative cell-associated HIV mRNA copy number was normalized to cell equivalents using 18S rRNA expression and the comparative Ct method (83).

### Linker-mediated (LM)-PCR and integration site data analysis

HIV-1 IS analysis was performed using LM-PCR and high-throughput sequencing as described previously (43). In brief, Timer populations from bulk-infected Jurkat cells with the HIV_Timer_ molecular construct were sorted and genomic DNA was extracted using the DNeasy Blood and Tissue Kit (QIAGEN) according to the manufacturer’s recommendations. About 2 µg of genomic DNA was sheared by sonication with a Picoruptor (Diagenode, S.A., Belgium) instrument to a size of 300–400 bp in length. DNA end was repaired and addition of adenosine at the 3’ end of the DNA was performed with NEBNext Ultra II End Repair/dA-Tailing Module (New England Biolabs, Cat# E7546). A linker was ligated to the ends of DNA using the NEBNext Ultra II Ligation Module (New England Biolabs, Cat# E7595). The junction between the 3’LTR of HIV-1 and host genomic DNA was amplified using primers targeting the 3’LTR and linker regions.

The first PCR amplicons were purified using a QIAquick PCR purification kit (QIAGEN), and then second PCR was performed. The following thermal cycler conditions were used for both PCRs: 96 °C for 30 s (1 cycle); 94 °C for 5 s, 72 °C for 1 min (7 cycles); 94 °C for 5 s, 68 °C for 1 min (13 cycles); 68° C 9 min (1 cycle) and hold at 4 °C. The second PCR amplicons were purified using a QIAquick PCR Purification Kit (QIAGEN), followed by Axy Prep Mag PCR cleanup bead purification (AXYGEN, Cat# MAG-PCR-CL-50). Purified PCR amplicons were quantified using the Agilent 4150 TapeStation system, and quantitative PCR was performed using primers P5 and P7 (GenNext NGS library quantification kit, Toyobo, Code# NLQ-101). LM-PCR libraries were sequenced using Illumina MiSeq paired-end reads, and the resulting FASTQ files were used for downstream analysis. Three FASTQ files, including Read1, Read2, and Index Read, were obtained from the IIIumina MiSeq. Read1 corresponded to sequencing data generated by a primer within the HIV-1 LTR region and Read2 to data generated by a primer within the linker. The Index Read corresponds to the 8-bp index sequence in the linker. First, we identified clusters on the flow cell with high sequencing quality of the Index Read (Phred quality score >30 at each position of the 8-bp index read) using an R code kindly provided by Michi Miura (Imperial College London, UK). Subsequently, we removed adaptor sequences from Read1 and Read2 selected reads with the LTR sequence in Read1 (TAGCA for HIV-1) and performed a cleaning step to remove reads that were too short or had a too low Phred score. The cleaned sequencing reads were aligned to the Homo sapiens genome assembly hg19 with HIV-1 (GenBank: K03455.1) using the BWA-MEM algorithm (84). We further processed the data to remove the unmapped reads, paired reads mapped to different chromosomes, and reads including 5’ LTR sequences. We then exported files containing information on the integration sites, DNA shear sites, and the number of reads. BED files were generated from exported files containing information on integration sites. RefSeq gene data were obtained from UCSC tables (https://genome-asia.ucsc.edu/). We analyzed the genome integration site preferences. The positions of the RefSeq genes were compared to the integration sites using the R package hiAnnotator (http://github.com/malnirav/hiAnnotator).

### DNA-capture-seq library and data analysis

HIV-1 DNA-capture-seq was performed as described previously (51) with minor modifications. Briefly, gDNA was extracted from Timer clones using the DNeasy Blood and Tissue Kit (Qiagen) according to the manufacturer’s instructions. 1 ug of genomic DNA was sheared by sonication using a Picorupter (Diagenode s.a., Liege, Belgium) to obtain fragments with an average size of 300 bp. Libraries for NGS were prepared using the NEBNext Ultra II DNA Library Prep Kit for Illumina (Cat# E7645) following the manufacturer’s instructions. To enrich the viral fragments contained in the synthesized libraries, several libraries were pooled to perform the capture step. The pooled libraries were mixed with virus-specific 161 biotinylated probes in the presence of human Cot-1 DNA (Invitrogen, Cat#15279011) and xGen Universal Blocking Oligos (IDT) for the hybridization step. A series of washing steps were performed using DNA xGen lockdown reagents (IDT), following the manufacturer’s recommendations. The quality of the enriched DNA libraries was evaluated and quantified by electrophoresis using the TapeStation 4150 system (Agilent Technologies). Finally, the multiplexed libraries were subjected to cluster generation using a MiSeq Reagent Kit v3 (150 cycles) in MiSeq sequencing systems (Illumina) with 2 x 75 bp reads. Three FASTQ files, Read1, Read2, and Index Read, were obtained from the Illumina MiSeq. We first performed a data-cleaning step using an in-house Perl script (kindly provided by Dr. Michi Miura, Imperial College London), which extracts reads with high Index Read sequencing quality (Phred score > 20 at each position of the 8-bp index read). Subsequently, we removed adaptor sequences from Read1 and Read2 followed by a cleaning step to remove reads with Phred scores that were too short or too low, as previously described (43). Cleaned sequencing reads were aligned to the reference genome using the BWA-MEM algorithm (84). To determine the complete sequence of the provirus in different cell lines and to infer its structure, the reference genome to which they were aligned included the entire human genome (hg19) and the complete HXB2 sequence as an independent chromosome. To properly determine the integration sites, the HXB2 sequence in the reference genome was included on two different chromosomes: viral LTRs (HIV_LTR) and proviral sequences without LTR (HIV_noLTR). We used SAMtools and Picard (https://github.com/broadinstitute/picard) for further data processing and clean-up, including the removal of reads with multiple alignments and duplicated reads. The Jurkat ChIP-seq datasets are described below. The final aligned files were visualized using Integrative Genomics Viewer (IGV 2.8.13).

### RNA-seq

For Jurkat Timer clones JO#10 and JO#19, RNA was extracted using the RNeasy Mini Kit (Qiagen) according to the manufacturer’s instructions with DNase I treatment. We ensured RNA quality using the Agilent Bioanalyzer 4150 TapeStation system. An RNA library was prepared using the NEBNext Ultra II directional RNA library prep kit for Illumina and sequenced using Illumina NextSeq 500. RNA sequences were processed using Cutadapt, PRINSEQ, STAR alignment, and SAMtools. Three independent biological replicates were analyzed for each clone. The Jurkat mRNA-seq dataset was obtained from the NCBI Sequence Read Archive (SRA) database under the GEO accession number GSM2171783 and processed in the same manner. The final aligned files were visualized using the UCSC Genome Browser (https://genome-asia.ucsc.edu)

### ChIP seq Datasets, epigenetic data analysis, and Hi-C

Chromatin data (ChIP-seq) from Jurkat uninfected T-cells was downloaded from Sequence Read Archive (SRA) data repository: H3K27ac (SRR12884765, SRR1603650, SRR1509753, SRR2043614), H3K4me3 (SRR577482, SRR577483), H3K4me1 (SRR7782877), H3K36me3 (SRR11783979, SRR11783980), H3K27me3 (SRR12884766, SRR647929, SRR11903008) and H3K9me3 (SRR13191912, SRR12884767). The FASTA files obtained were processed with in-house scripts using Cutadapt, PRINSEQ, BWA alignment, and SAMtools to obtain the mapped BAM files. Peak calling was performed using MACS2 and the corresponding input files for each histone mark. Features of histone modifications in the genomic environment flanking HIV-1 were analyzed using a Perl script (v5.22.0) as described previously (85). Hi-C for parent Jurkat T cells was obtained from NCBI GEO (GSE122958). Hi-C correlation matrices were generated using FAN-C (https://github.com/vaquerizaslab/fanc) and Cooler (https://github.com/mirnylab/cooler), at a resolution of 1 Mb.

### Statistical analysis

Statistical analyses were performed using GraphPad Prism 9 software (GraphPad Software, Inc., CA, USA). Fisher’s exact test was used to compare integration tendencies across Timer populations. Statistical significance was defined as *P* < 0.05.

## Acknowledgments

We are grateful to Y. Matsuoka, A. Murakawa, N. Monde, and H. Terasawa for technical and administrative support. This work was supported by research grants from the Japan Agency for Medical Research and Development (AMED) (JP23fk0410052, JP23wm0325068, JP23jm0210074, and JP23fk0410040) awarded to Y. S., a Biotechnology and Biological Sciences Research Council (BBSRC) David Phillips Fellowship (BB/J013951/2), a Medical Research Council (MRC) grant (MR/S000208/1) to M.O.. This work was supported by the JSPS Core-to-Core Program and JST MIRAI. The funders had no role in the study design, data collection, data interpretation, or discussions regarding submission for publication.

## Author Contributions

M.O. and Y.S. contributed to the study conceptualization. O.R. and Y.S. designed research; O.R., K. Monde, K.S., A.R., W.S., S.A.R. S.N.S, C.M., and H.T. performed research; K. Monde, M.O., and H.T. contributed new reagents/analytic tools; O.R., K.S., B.J.Y.T., K.N., K. Maeda, and Y.S. analyzed data; O.R. and Y.S. wrote original draft; All authors reviewed and revised the manuscript draft and approved the final version for submission.

