## Supplemental figures 1-8 for "HIV-Tocky system to visualize proviral expression dynamics"

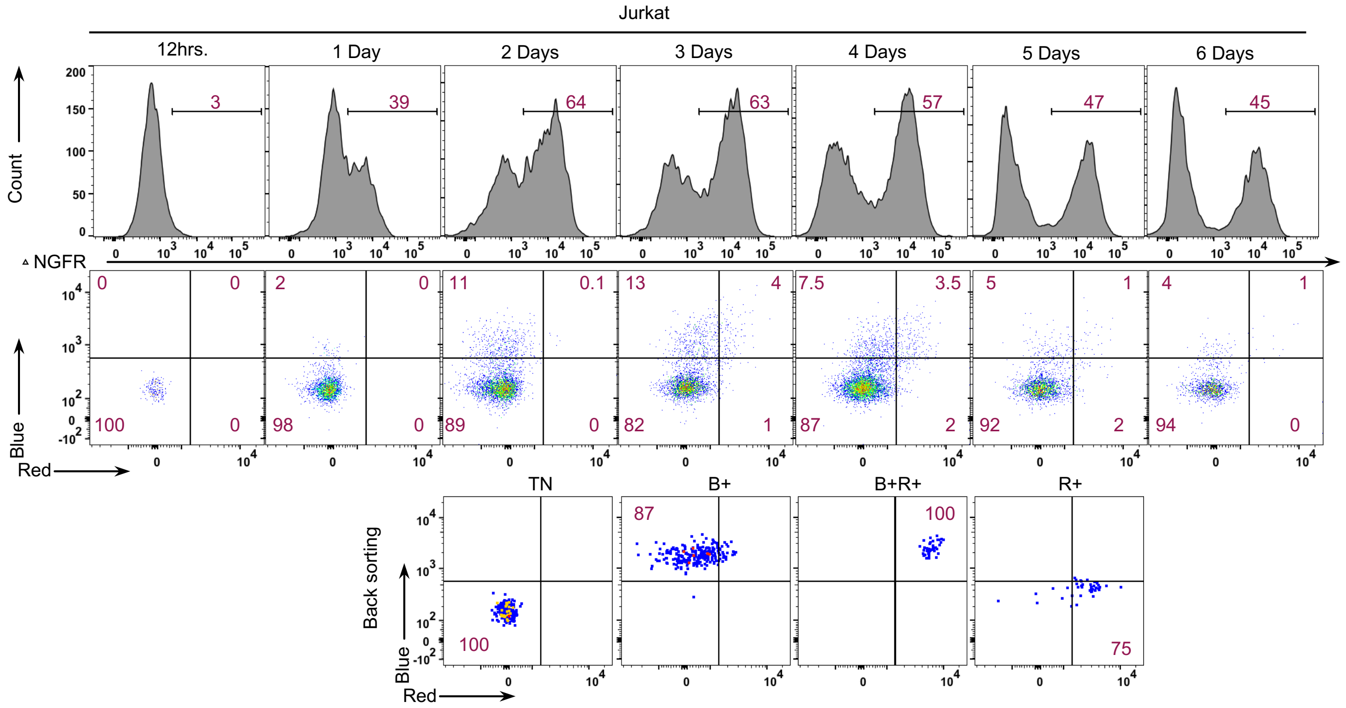


**Fig. S1. Jurkat infection and cell sorting for viral transcript quantification**
HIV$_{TNGFR}$ infection into Jurkat T cells 12 hrs. to 6 dpi; upper panel showing NGFR+ cells, middle panel showing Timer populations' distribution and lower panel showing purity check of sorted Timer populations (TN, B+, B+R+ and R+).

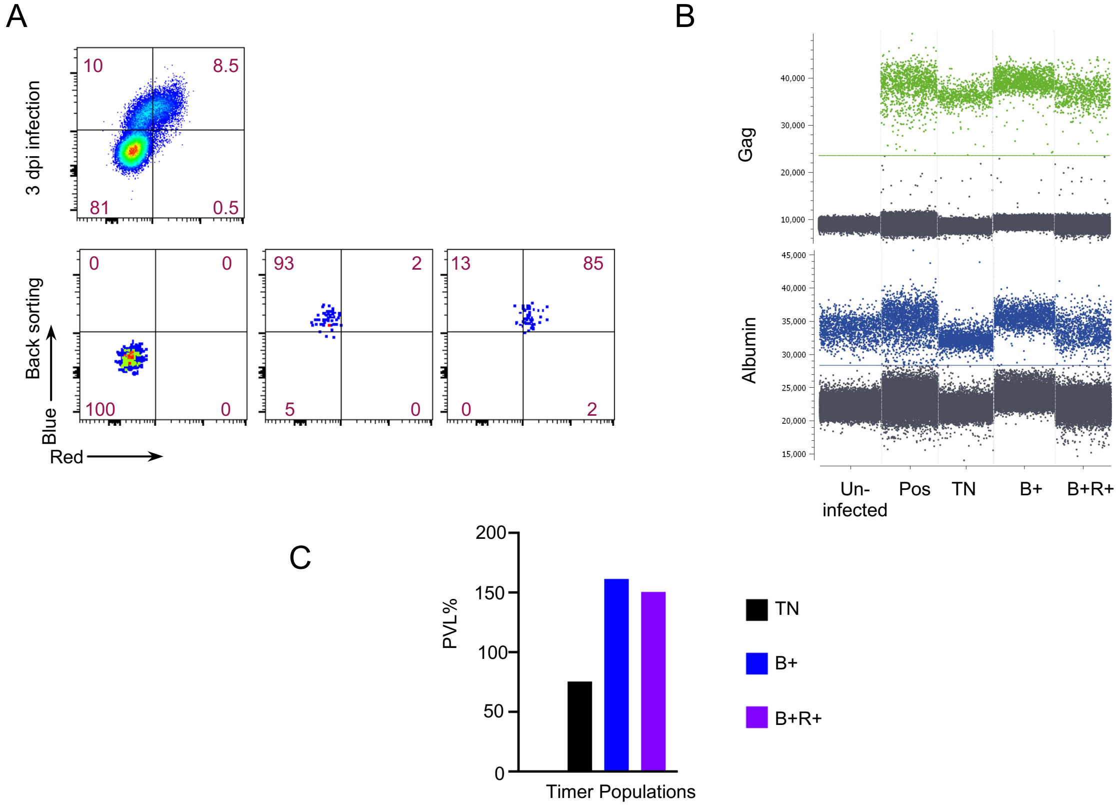


**Fig. S2. Proviral load calculation for Timer populations**
(A) HIV$_{Timer}$ infection into Jurkat T cells at 3 days post-infection used for sorting Timer fractions (TN, B+, and B+R+). (B) 1D dot plot showing droplet fluorescence intensity on the y-axis and each sample tested on the x-axis for *Gag* gene (upper panel) and *Albumin* gene (lower panel). (C) Proviral load % (PVL%) for each Timer population is calculated by the following formula (copy number of HIV-1 gag DNA)/[(copy number of Albumin)/2] x100.

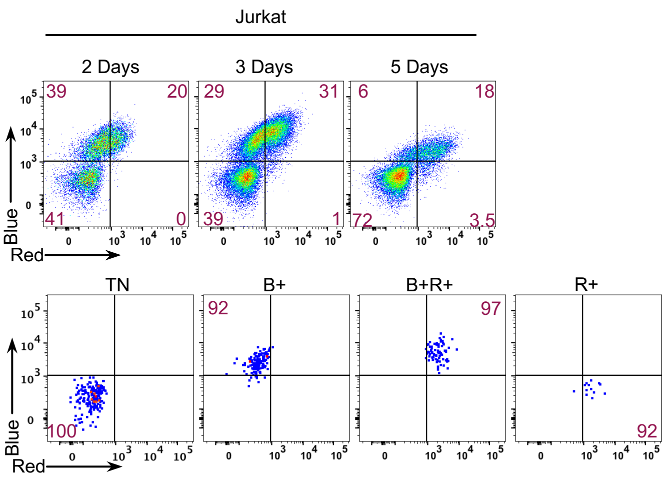


**Fig. S3. Jurkat infection and cell sorting for IS analysis**
$HIV_{Timer}$ infection into Jurkat T cells 2,3 and 5 dpi; upper panel showing Timer populations' distribution and lower panel showing purity check of sorted Timer populations (TN, B+, B+R+ and R+).

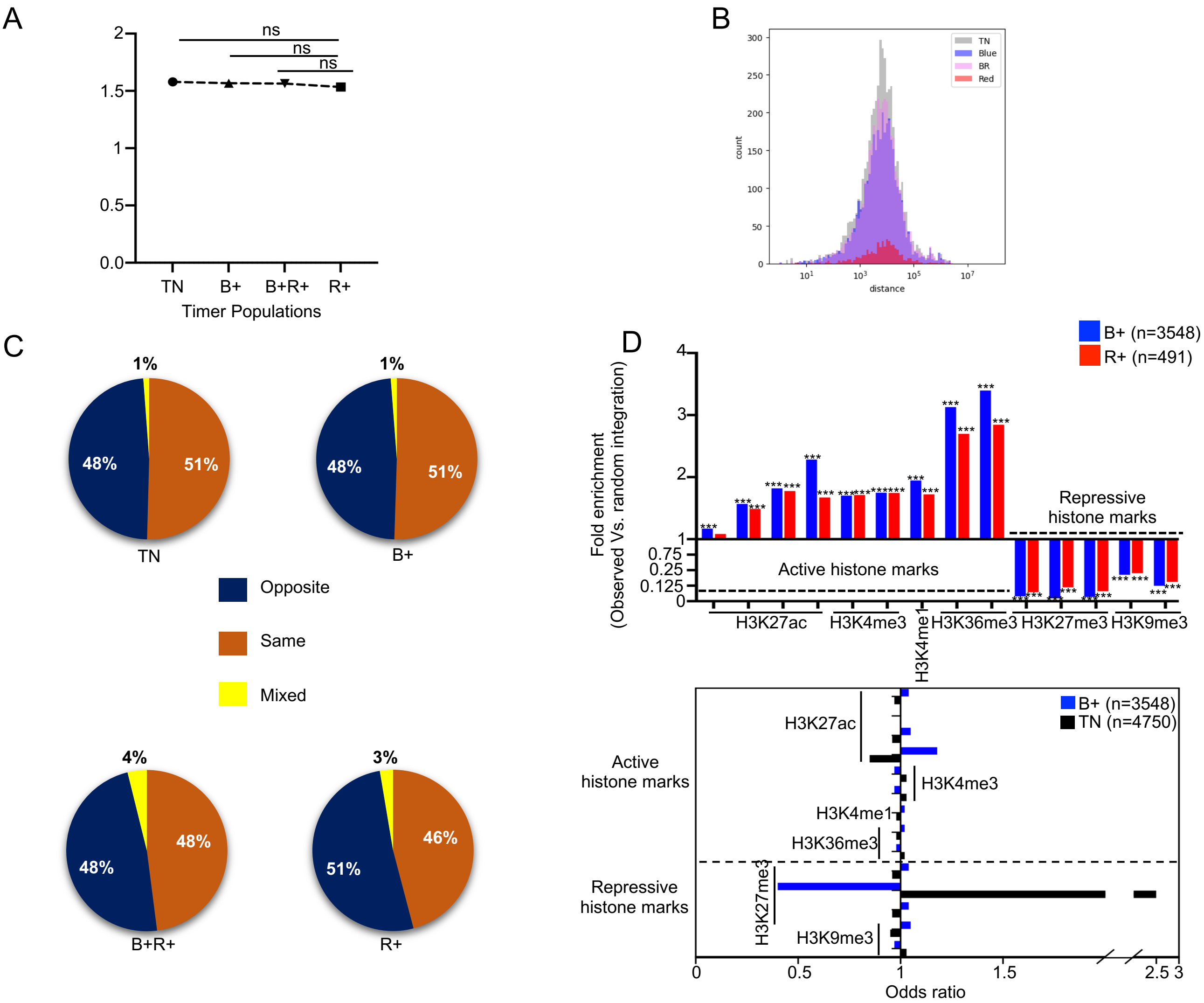


**Fig. S4. Integration site tendencies of various Timer populations**
(A) List of genes of integration from each Timer sorted population (TN, B+, B+R+, and R+) were checked for their basal (pre-integration) TPM values from Jurkat un-infected RNA-seq datasets. Mean $Log_{10}$ (TPM+1) values were calculated and plotted for each Timer population. Statistical significance was calculated by t-test between (TN and R+), (B+ and R+), and (B+R+ and R+). (B) Histogram showing the distribution of ISs within each Timer population in relation to chromosomal distance to the most proximal TSS. Statistical significance was calculated by Steel-Dwass's multiple comparison test. (C) Pie charts showing percentages of the same or opposite orientation of provirus integration relative to host gene in each Timer population. (D) Upper panel: Integration frequencies near the activating histone marks including H3K27ac, H3K4me3, H3Kme1, and H3K36me3, and the repressive histone marks with H3K27me3 and H3K9me3 comparing B+ to R+ populations. Integration sites within 2 kb of each histone mark were compared to random expected values. Fold enrichment is represented as the ratio of observed sites/random expected sites. Lower panel: The odds ratio of viral integration sites within ±2 kb of the same histone marks in the upper panel comparing TN and B+. Statistical significance was assessed by Fischer's exact test. *<.05, **<.01, ***<.001.

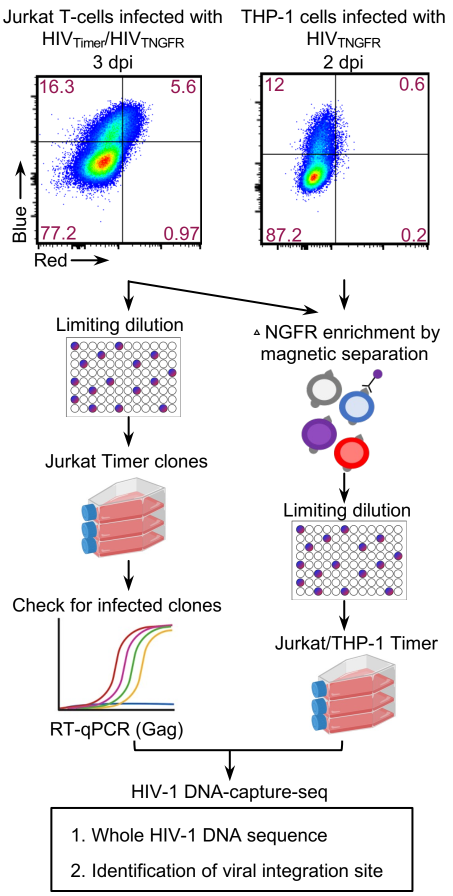


**Fig. S5. Establishment of HIV-Timer clones**
Experimental workflow of Timer clone generation with infected Jurkat T cells infection or THP-1 cells. Jurkat T cells (left side) were infected with $HIV_{Timer}$ and limiting dilution was performed at 3 dpi. qPCR check for infected clones was performed. Right side: THP-1 cells or Jurkat T cells were infected with $HIV_{TNGFR}$. Beads sorting for NGFR+ cells was performed at 3 dpi. After clones' propagation; DNA capture sequencing was performed.

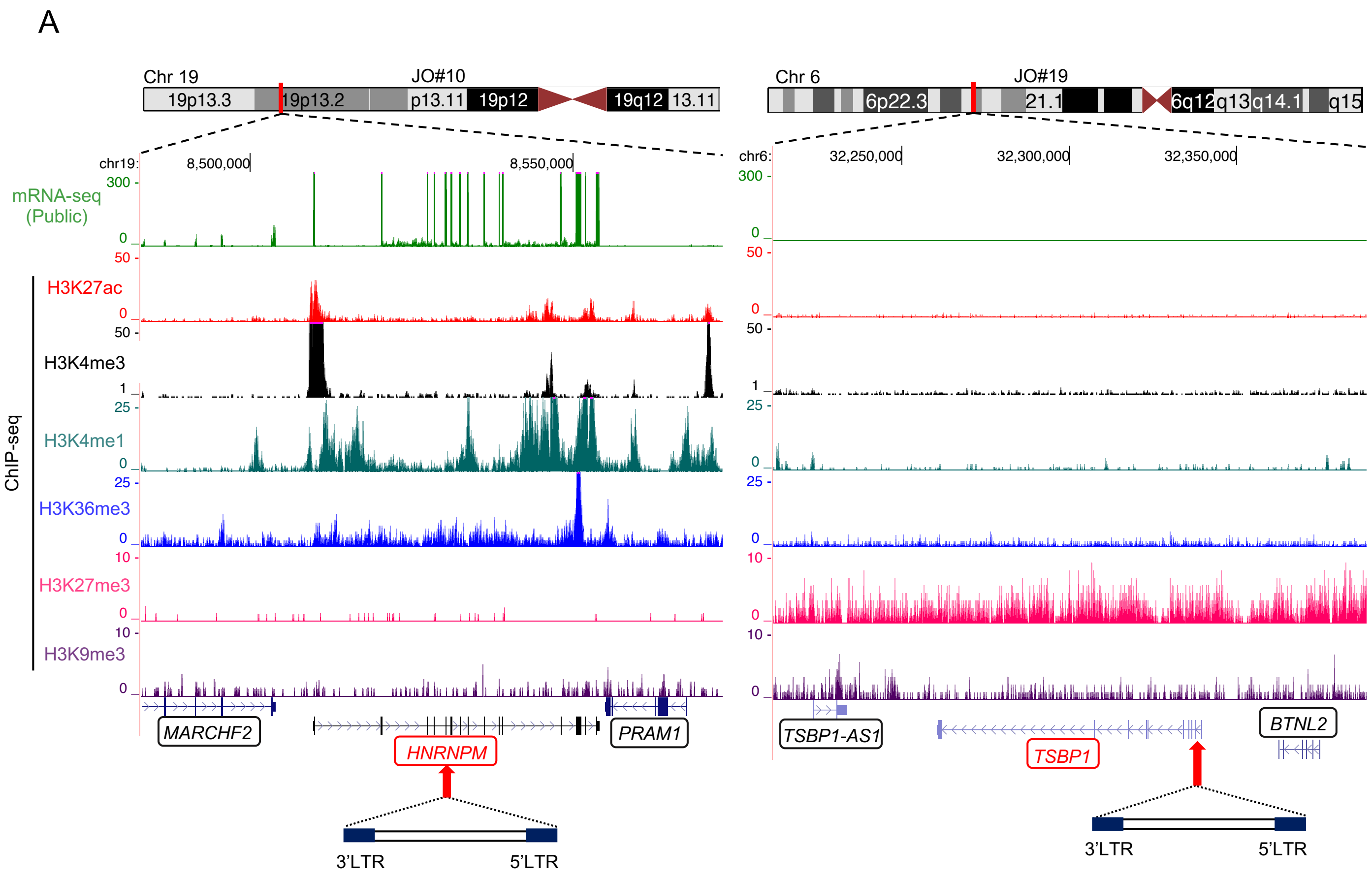


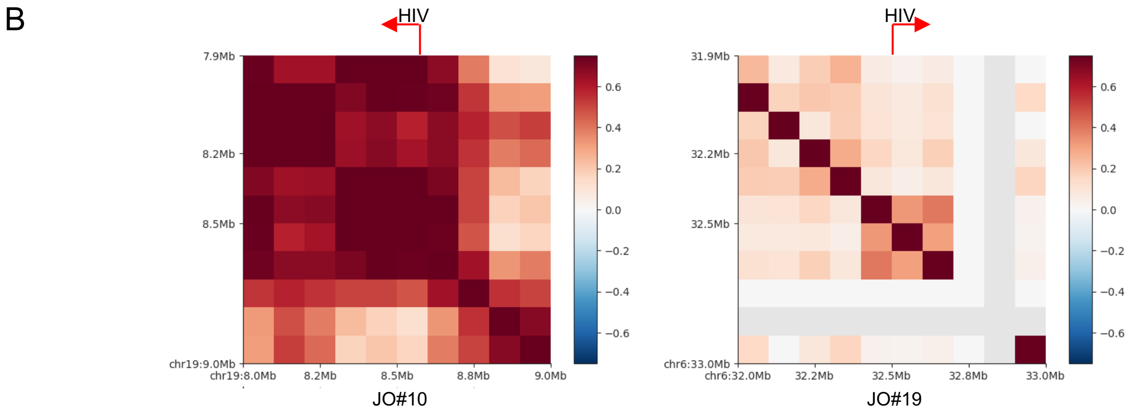


**Fig. S6. Integration environment of Timer Clones**
(A) RNA-seq data and ChIP-seq data sets of Jurkat T cells were blotted by UCSC Genome Browser (https://genome-asia.ucsc.edu) and shown in respect of integration genes of Timer clones JO#10 (left panel) and JO#19 (right Panel). Schematic for the position and directionality of provirus integration in each clone is demonstrated at the bottom. (B) Hi-C correlation matrices for Jurkat Timer clones 10 and 19: The correlation matrix illustrates the correlation (range from blue to red) between the intrachromosomal interaction profiles of every pair on 1-Mb loci along chromosome 19 (left panel) and chromosome 6 (right panel).

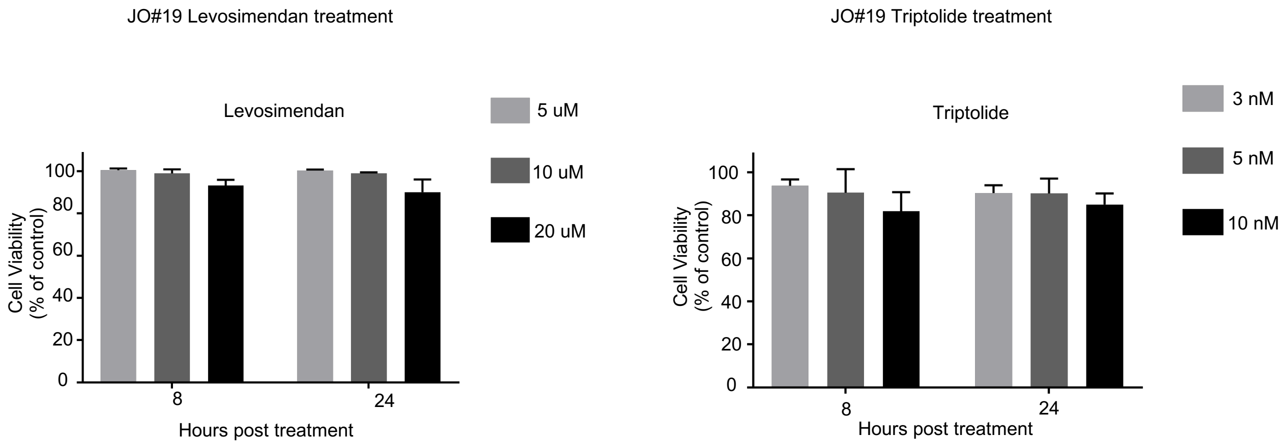


**Fig. S7. Cell viability assay using LPAs on JO#19**
Jurkat T cells or JO#19 were treated with Levosimendan, Triptolide, or DMSO control in shown concentrations for 8 or 24 hrs. At 8 or 24 24hrs, cells were treated with cell counting kit-8 reagent to check for cell viability. Cell viability % is calculated relative to values from un-infected Jurkat cells as control.

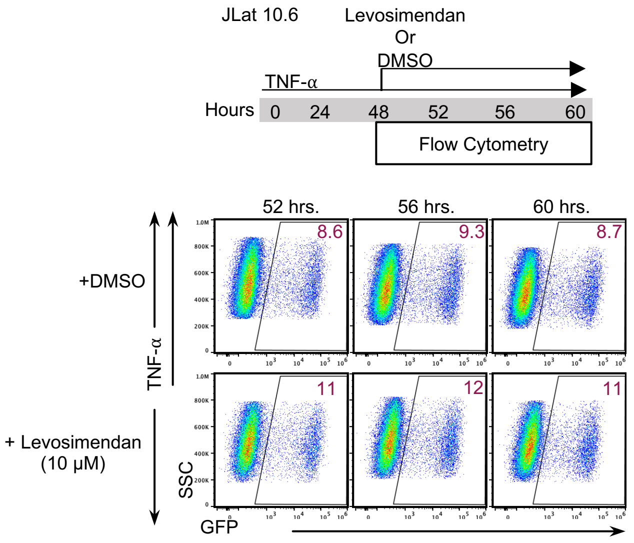


**Fig. S8. JLat 10.6 treatment with Levosimendan or DMSO**
JLat 10.6 cells were stimulated with TNF-α (10 ng/uL) for 48 hrs., an LPA drug or DMSO was further added for 12 hrs., and the changes in GFP expression were monitored. Flow plots demonstrating GFP transitions under Levosimendan (10 μM) treatment are shown.
